# Impact of infusion conditions and anesthesia on CSF tracer dynamics in mouse brain

**DOI:** 10.1101/2025.01.21.634133

**Authors:** Yuran Zhu, Junqing Zhu, Chenxin Ni, Anbang Chen, Longshun Li, Yue Gao, Andrew J. Shoffstall, Xin Yu

**Author notes:** Address correspondence to: Xin Yu, Sc.D., Wickenden 430 10900 Euclid Avenue Cleveland, OH 44106, USA.

## Abstract

Tracer imaging has been instrumental in mapping the brain’s solute transport pathways facilitated by cerebrospinal fluid (CSF) flow. However, the impact of tracer infusion parameters on CSF flow remains incompletely understood. This study evaluated the influence of infusion location, rate, and anesthetic regimens on tracer transport using dynamic contrast-enhanced MRI with Gd-DTPA as a CSF tracer. Infusion rate effects were assessed by administering Gd-DTPA into the cisterna magna (ICM) at two rates under isoflurane anesthesia. Anesthetic effects were evaluated by comparing transport patterns between isoflurane and ketamine/xylazine (K/X) anesthesia at the slower rate. Gd-DTPA transport was also examined after lateral ventricle (ICV) infusion, the primary site of CSF production. The results demonstrate that, besides anesthesia, both the location and rate of infusion substantially affected solute transport within the brain. ICV infusion led to rapid, extensive transport into deep brain regions, while slower ICM infusion resulted in more pronounced transport to dorsal brain regions. Cross-correlation and hierarchical clustering analyses of region-specific Gd-DTPA signal time courses revealed that ICM infusion facilitated transport along periarterial spaces, while ICV infusion favored transport across the ventricular-parenchymal interface. These findings underscore the importance of experimental conditions in influencing tracer kinetics and spatial distribution in the brain.

## Introduction

In vivo tracer imaging in preclinical models has been pivotal in evaluating solute transport in the brain, especially in advancing our understanding of the glymphatic system over the past decade ^1,2^. These studies typically involve cannulation and tracer infusion, mostly commonly into the cisterna magna (ICM), followed by dynamic imaging to monitor tracer transport and clearance within the brain. Of particular interest, dynamic contrast-enhanced MRI (DCE-MRI) provides the opportunity to track contrast agent-induced signal changes across the entire brain ^3,4^. The large field-of-view offered by DCE-MRI is crucial in understanding both the inflow of tracers along periarterial pathways and their outflow through multiple routes, including along perivenous spaces within the brain parenchyma, cranial and spinal nerves, and meningeal lymphatic vessels ^5^. DCE-MRI has been applied in studies ranging from elucidating the structure and function of CSF flow pathways to studying clearance dysfunctions under various pathophysiological conditions, including sleep-wake cycle ^6^, anesthesia ^7–9^, body postures ^10^, and cardiac function ^11^.

The rich spatial and temporal information provided by DCE-MRI also allows for a comprehensive assessment of solute transport characteristics using various image and statistical analysis tools ^12^. Voxel-wise clustering has been used to delineate CSF transport patterns and highlight shared transport characteristics within the same clusters^3^. Correlating solute transport patterns with anatomical structures provides additional insights into the architecture and functionality of the glymphatic system. To achieve this, atlas-based co-registration is essential for generating time courses of contrast agent-induced signal changes specific to individual brain regions, allowing for the depiction of solute transport kinetics across the brain aligned to their anatomical structures ^13,14^. This approach is particularly valuable for analyzing regions of pathophysiological interest, where modeling approaches can be integrated to probe potential alterations or dysfunctions in CSF transport behavior ^15^.

In this study, we present a comprehensive approach to examine the transport kinetics and spatial distribution of Gd-DTPA following its infusion into the CSF spaces of the mouse brain by DCE-MRI. This approach integrates maximal intensity projections, region-specific time-course analysis, and hierarchical clustering based on cross-correlation analysis. We investigated the transport kinetics and distribution patterns of Gd-DTPA in relation to the CSF production rate, primary CSF production site, and physiological factors influenced by anesthetic regimens. Our results demonstrate that besides anesthesia, both infusion location and infusion rate are key factors impacting the observed kinetics and distribution patterns of Gd-DTPA transport.

## Material and methods

### Animals

The animal protocol was approved by the Institutional Animal Care and Use Committee of Case Western Reserve University in accordance with the Guide for the Care and Use of Laboratory Animals and the Public Health Service Policy on Humane Care and Use of Laboratory Animals. The experiments were reported in compliance with Animal Research: Reporting in Vivo Experiments (ARRIVE) guidelines. All surgery and MRI scans were performed under isoflurane or ketamine/xylazine anesthesia, and all efforts were made to minimize suffering. The experiments were performed on 11- to 12- week-old male C57BL/6 mice (Jackson Laboratories, Bar Harbor, ME, US). The average body weight at the time of MRI scan was 27.5 g. The animals were housed in a temperature- and humidity-controlled environment with ad libitum access to food and water and a 12-hour light-dark cycle.

The animals were randomly assigned to one of the four experimental groups. All animals received a total of 10 µL of 12.5-mM Gd-DTPA diluted in artificial CSF (aCSF, Tocris BioScience, Minneapolis, MN, US). One group of mice received a relatively fast infusion rate of 1 µL/min, while the other three groups received a slower infusion rate of 0.33 µL/min. Infusion conditions included: 1) intra cisterna-magna (ICM) infusion at 1 µL/min for 10 min under isoflurane (ICM-ISO-fast, n=6); 2) ICM infusion at 0.33 µL/min for 30 min under isoflurane (ICM-ISO-slow, n=7); 3) ICM infusion at 0.33 µL/min for 30 min under K/X and low-dose isoflurane (ICM-K/X, n=6); and 4) intra-ventricular (ICV) infusion at 0.33 µL/min for 30 min under K/X and low-dose isoflurane (ICV-K/X, n=6).

### Anesthesia and cannulation procedures

All animals underwent cannulation surgery on a stereotaxis frame (Stoelting Co., Wood Dale, IL, USA) with the head secured by ear bars and a tooth bar. Body temperature was maintained at ∼37°C with a heating pad attached to the stereotaxic frame. Mice in ICM-ISO-fast and ICM-ISO-slow groups were first anesthetized with 2% isoflurane in an induction chamber and transferred to the stereotaxic frame. Anesthesia was maintained with 1 - 1.25% isoflurane delivered via a nose cone during the surgery and subsequent MRI scans. For mice in ICM-K/X and ICV-K/X groups, anesthesia was induced with an intraperitoneal (IP) injection of ketamine (75 mg/kg) and xylazine (5 mg/kg). Low-dose isoflurane (0.5 - 0.75%) was administered as supplemental anesthesia during the MRI scans. For mice undergoing ICM cannulation, a polyethylene micro-catheter (0.13 mm ID × 0.25 mm OD, Scientific Commodities, Lake Havasu City, AZ, US) was inserted into the cisterna magna as previously described ^14,16^. Mice in ICV-K/X groups underwent cannulation into the right lateral ventricle. Specifically, after placing the mouse on the stereotaxic apparatus, a vertical incision was made along the midline of the head. With the bregma and lambda aligned on the same horizontal plane, a craniotomy was performed with a drill directly above the right lateral ventricle. A silica capillary catheter (0.15 mm ID × 0.36 mm OD, Molex, Lisle, IL, US) was then cannulated at the coordinates: 0.3 mm caudal, 1.0 mm lateral, and 2.3 mm ventral to the bregma. Following the cannulation, surgical glue (VetBond, 3M, St. Paul, MN, US) and dental cement (Co-Oral-Ite Dental Mfg. Co., Diamond Springs, CA, US) were applied to secure the cannula for all animals.

### MRI protocol

All MRI studies were performed on a horizontal bore 9.4T Bruker BioSpin scanner operating on the ParaVision 360 v3.3 platform (Bruker BioSpin, Billerica, MA, US). All MRI acquisitions used an 86-mm volume coil for transmitting and a four-channel array coil for receiving. After the cannulation, the animal was transferred to an MRI-compatible cradle and placed in a prone position. Body temperature and respiration rate were monitored during the MRI scans. The body temperature was maintained at ∼37°C by blowing warm air into the scanner through a feedback control system (SA Instruments, Stony Brook, NY, US). The respiration rate was maintained at 90 - 110 breaths per minute (BPM) by adjusting the isoflurane dosage.

Dynamic changes induced by Gd-DTPA in the whole brain were tracked using a T1-weighted fast low angle shot (FLASH) sequence with the following parameters: TR/TE, 50/2.8 ms; flip angle, 20°; field of view (FOV), 20 × 16 × 14 mm^3^; matrix size, 150 × 120 × 105; yielding an isotropic resolution of 133 µm and a temporal resolution of ∼5 min. Imaging included a 2-average baseline scan followed by 25 single-average dynamic scans acquired before, during, and after Gd-DTPA infusion, covering a total imaging duration of 130 min.

### MRI image analysis

All image reconstruction and data analyses were performed using either in-house developed or open-source software in MATLAB (MathWorks, Natick, MA, US) or Python (Python Software Foundation, v.3.0). All image registration used an open-source toolbox, the Advanced Normalization Tools ^17,18,18,19^. All dynamic images were motion-corrected by co-registration to their respective baseline images through affine transformation. For spatial standardization, a representative animal was selected from each group, and its baseline image was co-registered to an MRI mouse brain atlas by affine and deformable transformations ^20^, enabling segmentation of the whole brain into 20 regions of interest (ROIs). Subsequently, images from the remaining animals in each group were then registered to that of their corresponding representative animal’s image.

Maximum intensity projection (MIP) maps were generated by performing voxel-wise searches for maximal signal intensity across the dynamic image series. Using the 20 ROIs generated from co-registration with the brain atlas, the mean signal intensity in each ROI was calculated for each animal. The baseline signal was subtracted from the entire dynamic time series. Signal intensity in each ROI was then normalized by the maximal signal intensity from a small volume surrounding the infusion site, thereby eliminating inter-subject variations in T1-weighed signal intensity.

### Correlation and clustering analysis

Time-lagged cross-correlation analysis was performed to determine the maximal cross-correlation (mCC) and the corresponding lag time between two dynamic time series of signal intensity changes ^21^. Hierarchical clustering analysis was then applied to further evaluate transport kinetic correlations ^22^, with dissimilarity defined as 1 − *mCC* to quantify the distance between two ROIs. A dendrogram was generated with complete linkage to represent the hierarchical structure of the mCC matrix ^23–26^. Bootstrap analysis was performed with 1000 replications to evaluate the robustness of the clustering results ^27^. The uncertainty in clustering was evaluated using the approximately unbiased (AU) p-values and the bootstrap probability (BP) values. For mice in ICM-ISO-fast and ICM-ISO- slow groups, which showed similar signal intensity in most brain regions, mCC and the corresponding lag time were calculated in the same brain regions across the group to assess differences in transport kinetics under different ICM infusion rates.

## Results

### Distribution of Gd-DTPA in the whole brain

Figure 1 displays group-averaged dynamic images. Under isoflurane anesthesia, the slower ICM infusion rate of 0.33 μL/min promoted Gd-DTPA transport along the dorsal brain surface, whereas a faster rate of 1 μL/min favored transport along the ventral surface. The K/X anesthesia significantly enhanced Gd-DTPA transport along the ventral surface arteries and into the olfactory bulbs via the olfactory artery. In contrast, ICV infusion resulted in Gd-DTPA transport predominantly across the ventricular-parenchymal interface into deep brain regions, with delayed transport along the ventral surface.

**Figure 1.**
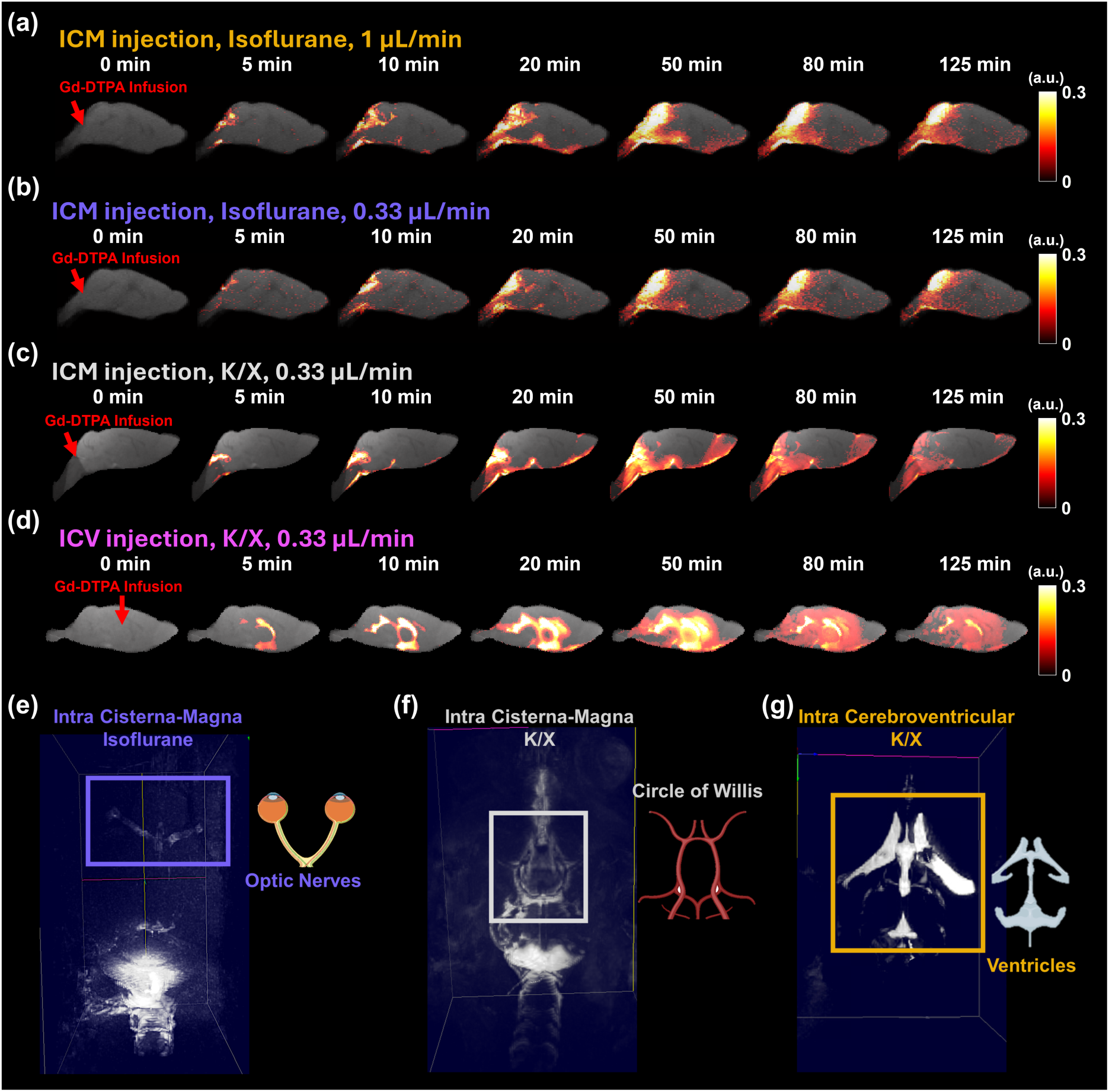
Contrast agent distribution at selected time points (a-d) and maximal intensity projection maps of Gd-DTPA distribution (e-g). Representative sagittal views of group- averaged images are overlaid with signal changes from the baseline. (a) Intra-cisterna- magna (ICM) infusion at 1 μL/min, under isoflurane anesthesia. (b) ICM infusion at 0.33 μL/min, under isoflurane anesthesia. (c) ICM infusion at 0.33 μL/min, under ketamine/xylazine (K/X) anesthesia. (d) Intracerebroventricular (ICV) infusion at 0.33 μL/min, under K/X anesthesia. (e) A representative mouse that received intra cisterna- magna (ICM) Gd-DTPA infusion at 0.33 μL/min under isoflurane anesthesia. The blue box indicates the optic nerves. (f) A representative mouse that received ICM Gd-DTPA infusion at 0.33 μL/min under ketamine/xylazine (K/X) anesthesia. The gray box indicates the circle of Willis and its periarterial space. (g) A representative mouse that received intra-cerebroventricular infusion at 0.33 μL/min under K/X anesthesia.

The MIPs of the entire brain more clearly depicted transport patterns in non-cortical regions (Figure 2). Under isoflurane anesthesia, Gd-DTPA transport from CM is relatively more prominent along the optic nerves, while K/X anesthesia enhances transport along the perivascular spaces in the circle of Willis. Gd-DTPA transport from the lateral ventricles showed significantly reduced distribution along major arteries.

**Figure 2.**
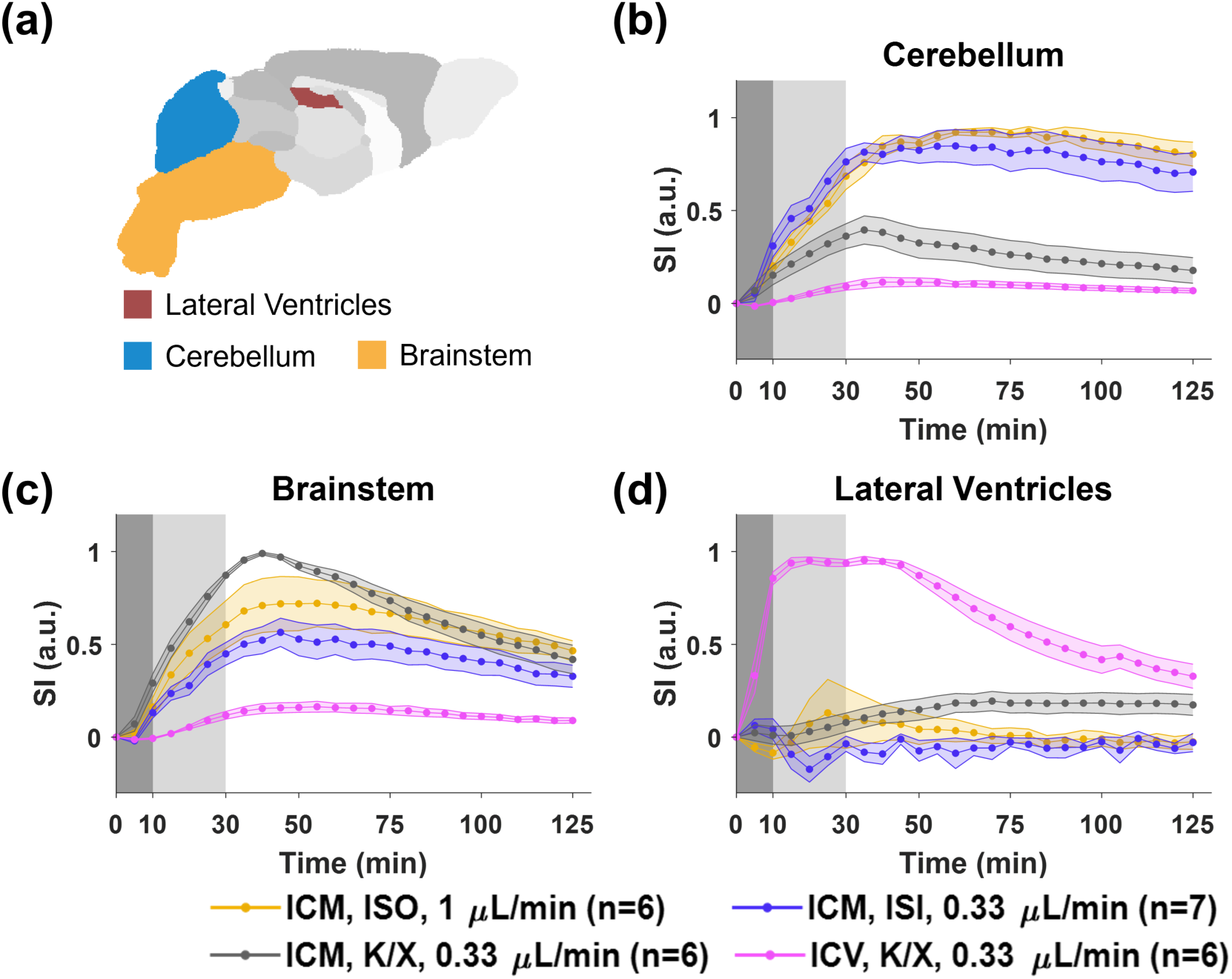
Dynamics of contrast agent transport at or proximal to infusion sites. (a) Segmentation of selected ROIs. (b)-(d) Time courses of signal changes in the selected ROIs. Gray bands indicate the time period of contrast agent infusion. Gray, magenta, yellow, and blue lines represent the mean time courses of signal changes. Shaded areas represent standard errors. ICM, intra-cisterna-magna infusion; ICV, intracerebroventricular infusion; ISO, isoflurane anesthesia; K/X, ketamine/xylazine anesthesia.

### Time-course signal changes

Time-course signal analyses revealed distinct kinetic differences among the groups (Figures 3-6, Figure 1s). In the three ICM groups, mice under isoflurane anesthesia exhibited persistent accumulation and slower washout near the infusion sites, regardless of infusion rate (Figures 3b-c). In contrast, mice under K/X anesthesia exhibited rapid Gd-DTPA exit via the brainstem (Figure 3c). The ICV group quickly reached signal saturation in the lateral ventricles during infusion, followed by a steady washout (Figure 3d).

**Figure 3.**
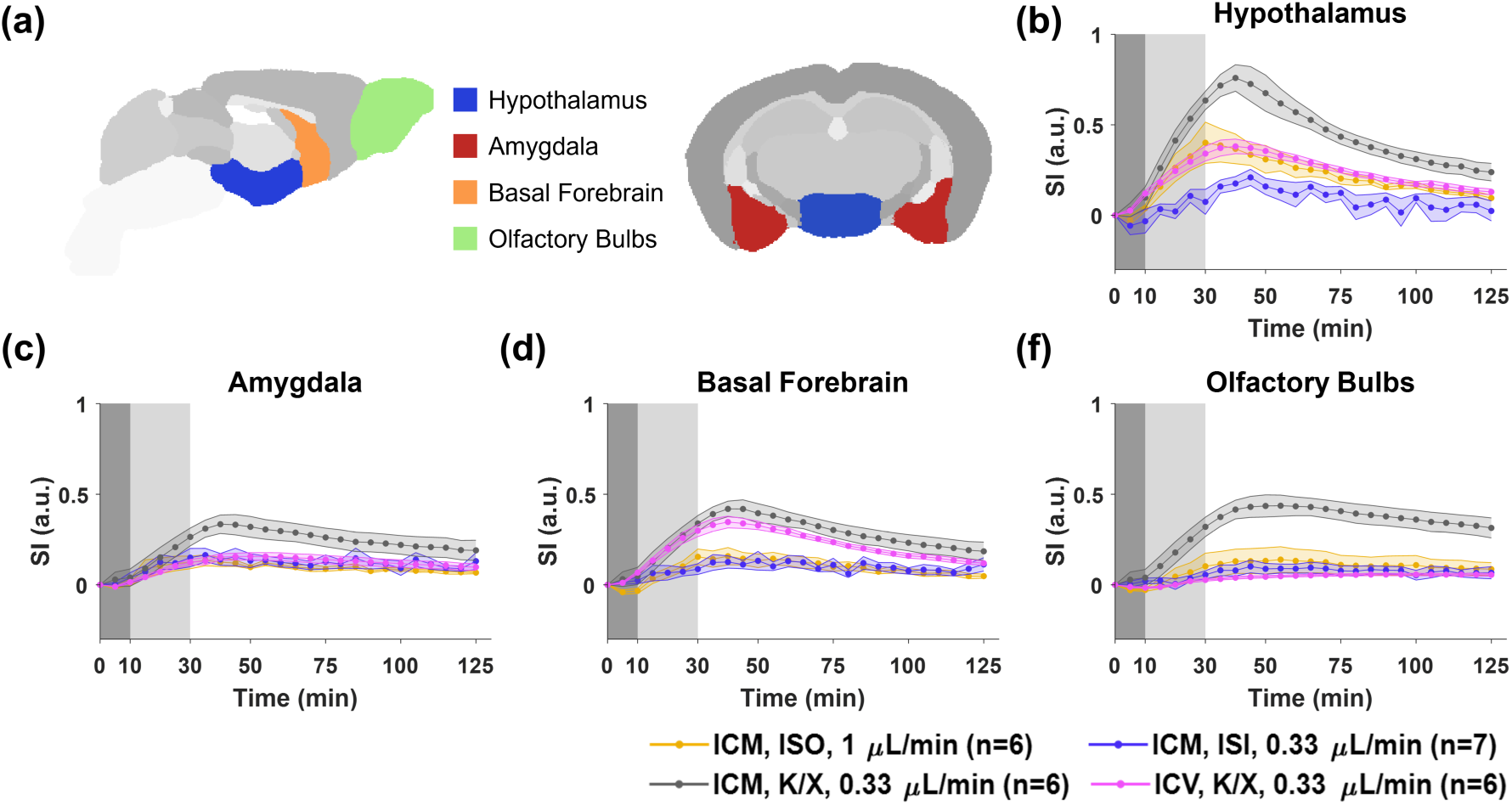
Dynamics of contrast agent transport in ventral brain regions. (a) Segmentation of selected ROIs. (b)-(f) Time courses of signal changes in the selected ROIs. Gray bands indicate the time period of contrast agent infusion. Gray, magenta, yellow, and blue lines represent the mean time courses of signal changes. Shaded areas represent standard errors. ICM, intra-cisterna-magna infusion; ICV, intracerebroventricular infusion; ISO, isoflurane anesthesia; K/X, ketamine/xylazine anesthesia.

Along the ventral surface, in regions closely associated with the major arteries of the brain, the ICM-K/X group exhibited the most significant Gd-DTPA transport, whereas the ICM-ISO-slow group showed the least significant transport (Figure 4). Notably, Gd-DTPA transport from the lateral ventricle (ICV infusion) to these regions differed drastically from the three ICM groups. The magnitude of signal change in the ICV-K/X group was comparable to that of the ICM-ISO-fast group in the hypothalamus (Figure 4b) and similar to both isoflurane groups in the amygdala (Figure 4c), and it was the lowest in the olfactory bulbs (Figure 4f). However, in basal forebrain (Figure 4d), an ROI adjacent to third ventricle, the ICV-K/X group showed comparable signal enhancement and transport kinetics to the ICM-K/X group.

**Figure 4.**
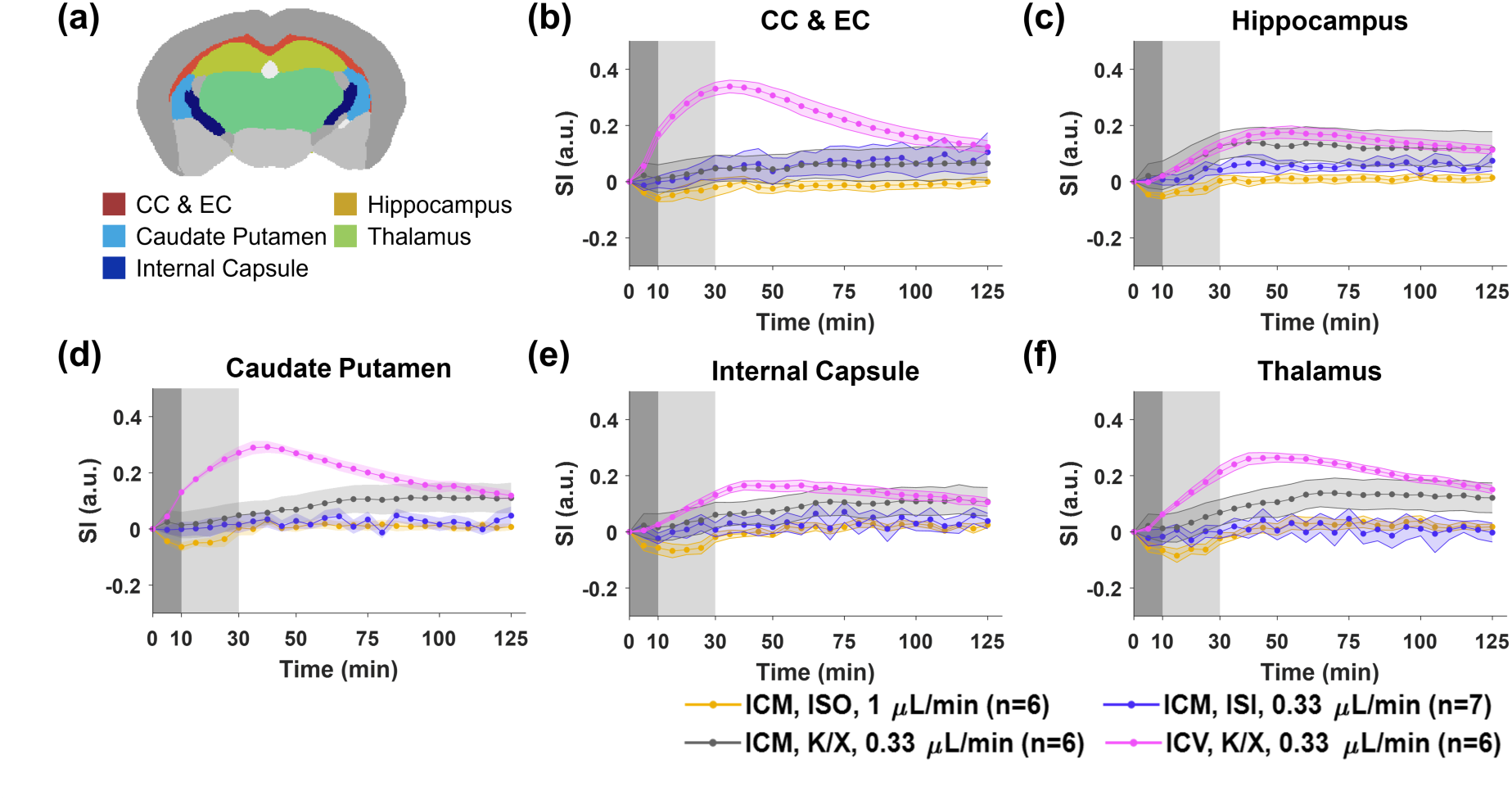
Dynamics of contrast agent transport in deep brain regions. (a) Segmentation of selected ROIs. (b)-(f) Time courses of signal changes in the selected ROIs. Gray bands indicate the time period of contrast agent infusion. Gray, magenta, yellow, and blue lines represent the mean time courses of signal changes. Shaded areas represent standard errors. ICM, intra-cisterna-magna infusion; ICV, intracerebroventricular infusion; ISO, isoflurane anesthesia; K/X, ketamine/xylazine anesthesia.

The time courses in the deep brain and dorsal surface regions are shown in Figs. 5 and 6. The ICV-K/X group showed extensive and rapid transport to the deep brain regions (Figure 5), with the highest and most rapid signal intensity changes in the corpus callosum and external capsule (CC & EC, Figure 5b), a region in close proximity to the lateral ventricles. The ICM-K/X group demonstrated more pronounced Gd-DTPA transport into the deep brain region compared to the two isoflurane groups. In contrast, the ISO groups exhibited more rapid and extensive Gd-DTPA transport into inferior colliculi, a posterior brain region on the dorsal brain surface (Figure 6c), while the neighboring superior colliculi did not (Figure 6d). All four groups showed limited signal changes in neocortex, particularly in the ICM-ISO-fast group (Figure 6b). These findings emphasize varying transport pathways based on infusion conditions and anesthesia.

**Figure 5.**
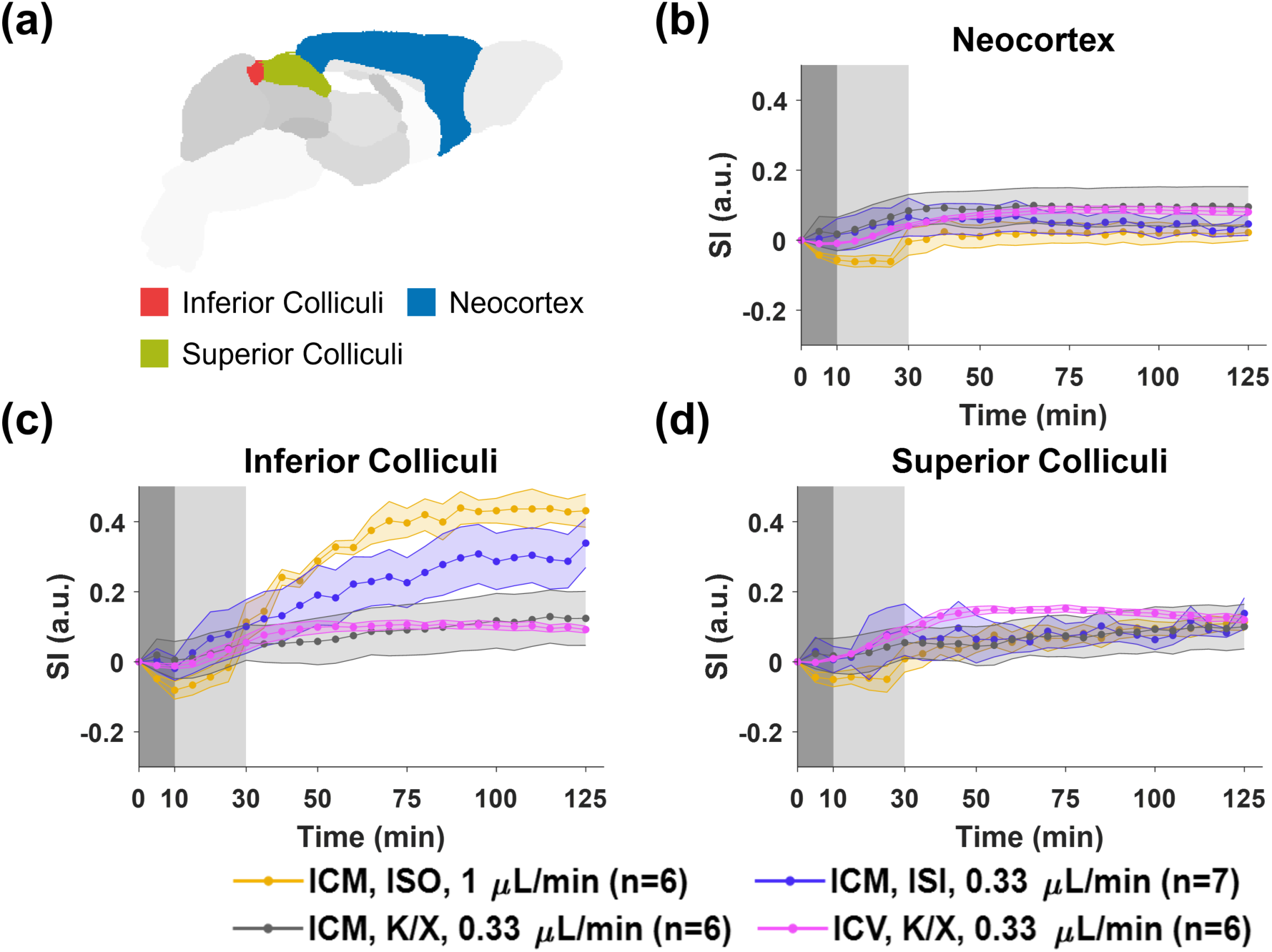
Dynamics of contrast agent transport in dorsal brain regions. (a) Segmentation of selected ROIs. (b)-(d) Time courses of signal changes in the selected ROIs. Gray bands indicate the time period of contrast agent infusion. Gray, magenta, yellow, and blue lines represent the mean time courses of signal changes. Shaded areas represent standard errors. ICM, intra-cisterna-magna infusion; ICV, intracerebroventricular infusion; ISO, isoflurane anesthesia; K/X, ketamine/xylazine anesthesia.

### Time-lagged cross-correlation analysis

The time courses from ICM-ISO-fast and ICM-ISO-slow groups were compared against each other by time-lagged cross-correlation analysis (Figure 7). The maximal cross-correlation coefficient (mCC) is a measure of the similarity in transport kinetic profiles between the two groups and the lag time indicates the relative delay in Gd-DTPA transport between the two groups. In the ventral brain regions, the mCC was above 90% in all ROIs, with minimal lag time except in the hypothalamus. This suggests that Gd-DTPA transport kinetics were largely similar between the two groups in these regions. In contrast, in the dorsal regions, the mCC decreased from 99% in the inferior colliculi to <50% in neocortex, suggesting significant dispersion in transport kinetics within the neocortical region between the two groups. Furthermore, the negative lag times indicate slower transport kinetics in the ICM-ISO-fast group compared to the ICM-ISO-slow group.

### Hierarchical clustering of brain regions

Figure 8 shows the comparison of potential transport pathways between the ICV- K/X and ICM-K/X groups by correlation-matrix-based hierarchical clustering analysis. A large mCC value between two ROIs suggests a high similarity in the transport kinetic profiles and the lag time indicates the relative delay in Gd-DTPA transport between the two regions. The clustering analysis identified three clusters with a dissimilarity value of < 0.4 for both groups. Bootstrap analysis with 1000 replications returned an approximately unbiased (AU) p-value of 0.93-1 for all clusters (Figures 8a&d). Figures 8b and 8c show the signal changes in each cluster were plotted against the individual ROIs in the same cluster.

For the ICV-K/X group, the lateral ventricles and their immediately adjacent regions were classified into the first cluster (C1), with AU p-values and bootstrap probabilities (BP) consistently at 1 (Figure 8a). Notably, the hypothalamus and basal forebrain, despite their proximity to large arteries on the ventral surface, were also included in this cluster. The second cluster (C2) predominantly comprised regions located on the posterior and ventral sides of the brain, while the third cluster (C3) encompassed dorsal brain regions on the anterior side. The ICM-K/X group demonstrated a markedly different Gd-DTPA transport pattern compared to the ICV-K/X group (Figure 8d). The C1 cluster included ROIs spatially proximal to the circle of Willis, whereas C2 contained brain regions further away from the major arteries. The majority of deep brain regions were grouped into the third cluster (C3) that shared much less similarity than the first two clusters.

## Discussion

This study investigated the impact of infusion site, infusion rate, and anesthesia on Gd-DTPA transport using DCE-MRI. Methods of ROI-based time course analyses were developed to identify potential transport pathways under varying experimental conditions. Consistent with previous observations in rodents ^7–9^, K/X anesthesia significantly enhanced periarterial Gd-DTPA transport compared to isoflurane. Notably, a slower infusion rate (0.33 μL/min) under isoflurane anesthesia resulted in significantly increased transport to dorsal brain regions. Furthermore, K/X promoted transport along major arteries, while isoflurane resulted in more pronounced transport along the optic nerves. The spatial distribution and kinetics of Gd-DTPA transport varied markedly between ICV and ICM infusion. In the ICV-K/X group, Gd-DTPA transport across the ventricular- parenchymal interface into deep brain regions was observed, preceding its transport to the ventral and dorsal brain surfaces. Time-lagged cross-correlation and hierarchical clustering analyses provided further insight into these distinctive transport pathways that reflected the influence of infusion sites.

The extensive spatial and temporal data provided by DCE-MRI enables a detailed assessment of solute transport in CSF and its correlation to various physiological or experimental factors ^3,12^. In this study, we focused on experimentally controllable parameters, including tracer infusion site, infusion rate, and anesthesia type. Image analysis reveals dynamic CSF transport patterns, capturing contrast agent transport not only within the brain but also along the optic nerves and cerebral arteries (Figure 2). Tracer transport in a specific region can be assessed through ROI-based analysis ^9^. Furthermore, brain-region-specific signal changes can be quantified by co-registering dynamic images with a brain atlas to compare differences across animal groups and brain regions ^13,14^. While direct observations of the magnitude and kinetics of signal changes are informative, additional statistical, clustering, and modeling analyses offer further insights into the CSF transport dynamics ^28^. Recent advancements include k-means clustering, optimal mass transport modeling, and compartment-modeling methods ^4,29–31^. Using a correlation-matrix-based hierarchical clustering method, this study compared potential Gd-DTPA transport patterns via different infusion routes. The analysis identified a strong transport correlation between the deep brain regions and a part of the ventral areas, including the hypothalamus and basal forebrain, with the lateral ventricles (C1, Figures 8a-b). In contrast, Gd-DTPA transport via ICM infusion displayed a preference for pathways along the major periarterial spaces of the ventral brain surface (Figures 8c-d).

Most DCE-MRI studies in animals have been conducted under anesthesia, with varying effects on observed solute transport patterns depending on the anesthetic regimens used ^3^. Recent research suggests that isoflurane may suppress CSF flow, whereas K/X or dexmedetomidine supplemented with low-dose isoflurane enhances solute transport ^7,9,11^. This is attributed to the vasoconstrictive effect of K/X on α2- adrenergic receptors in cerebral arteries, as opposed to the vasodilatory effects of isoflurane ^7,32^. Consistent with these findings, our study observed enhanced Gd-DPTA transport along major cerebral arteries under K/X anesthesia (Figure 2b), supported by time-course analyses of signal changes in ventral brain surface regions (Figure 4). In contrast, isoflurane-anesthetized animals showed more pronounced signal enhancement in the optic nerves compared to cortical and anterior ventral brain regions (Figure 2a). While optic nerves were not the primary focus of this study, they are considered crucial for CSF outflow. Historically, cranial and spinal nerves are estimated to drain 10-15% of CSF into the systemic lymphatic system via their perineural space ^33^. In mice, the ocular CSF route is particularly significant and may play a key role in CSF outflow ^34^. Recent findings suggest the existence of an ocular glymphatic system analogous to the brain’s^35^. A study by Tong and colleagues demonstrated that the accumulation of ICM-infused CSF tracers in the optic nerves is highly sensitive to infusion conditions, including tracer molecular size, infusion volume, and infusion rate ^36^. The current study extended these findings by examining the impact of anesthetic regimens. Isoflurane-anesthetized animals displayed more pronounced Gd-DTPA transport along the optic nerves compared to K/X-anesthetized animals under identical infusion conditions. Further investigations are warranted to integrate structural imaging of the optic nerves to explore the ocular CSF flow pathway further.

CSF production is a key factor influencing solute transport within the brain, yet it remains underexplored. It is generally accepted that CSF is primarily produced by the choroid plexus in the brain’s ventricles, from which it flows into the subarachnoid space surrounding the brain and spinal cord ^37,38^. Additional CSF may be produced via fluid secretion across capillary walls ^39^. Solute transport is likely driven by a pressure gradient between the production and drainage sites, with slightly higher pressure in the lateral ventricles ^33^. ICV infusion thus provides a valuable approach for investigating solute transport in relation to these natural CSF production sites. In the current study, substantial differences were observed in the spatial distribution and transport dynamics of Gd-DTPA based on infusion routes. These differences were evident in dynamic imaging (Figures 1c-d), brain-region-specific time courses (Figures 4-6, Figure S1), and clustering analyses (Figure 8). Consistent with findings in nonhuman primates, ICV infusion resulted in rapid and extensive Gd-DTPA transport to deep brain regions adjacent to lateral ventricles, while the ventral brain surface showed reduced signal enhancement and slower transport, despite their proximity to major cerebral arteries and the vasoconstriction effects of K/X (Figure 4). Furthermore, the strong transport correlations identified by clustering analysis between the lateral ventricles and deep brain regions in the ICV-K/X group suggest the involvement of the ventricular-parenchymal interface in CSF transport in addition to its circulation through the subarachnoid and periarterial spaces.

**Figure 6.**
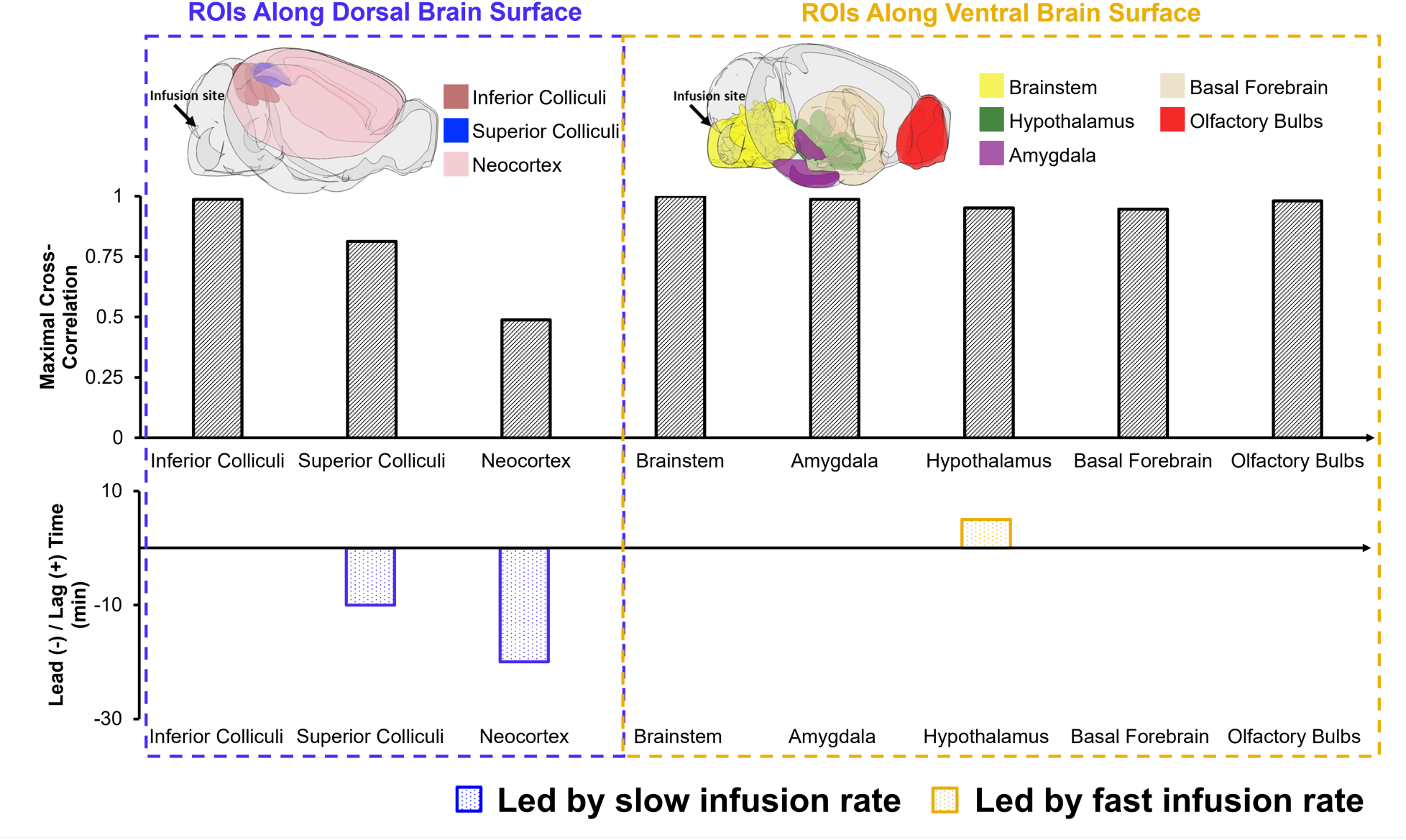
Comparison of Gd-DTPA transport kinetics at different infusion rates (ICM-ISO- fast vs ICM-ISO-slow) by time-lagged cross-correlation analysis. mCC: maximal cross- correlation coefficients. Negative lag time indicates faster transport kinetics in the ICM- ISO-slow group, whereas positive lag time indicates faster transport kinetics in the ICM- ISO-fast group.

While ICV infusion offers valuable insights, intracisternal magna (ICM) infusion remains the most widely used method for administering CSF tracers ^3^. Most studies involving mouse models have used an infusion rate of 1 μL/min, which is considerably higher than the natural CSF production rate of approximately 0.35 μL/min measured in the lateral ventricles ^40,41^. Prior research has demonstrated a 2.5-mmHg elevation in intracranial pressure (ICP) in mice receiving ICM infusion at 2 μL/min ^42^. Another study reported baseline ICP values of ∼4 mmHg in both isoflurane- and K/X-anesthetized mice^43^. These findings suggest that an infusion rate of 1 μL/min could induce a transient increase in ICP above its baseline level. In this study, two different ICM infusion rates were compared. Consistent with previous findings, Gd-DTPA transport into cortical regions on the dorsal brain surface was hindered at the faster rate of 1 μL/min. Conversely, the slower rate of 0.33 μL/min facilitated Gd-DTPA transport to these regions despite the use of isoflurane anesthesia, as depicted in Figure 6b. Time-lagged cross-correlation analysis revealed high correlations in Gd-DTPA transport kinetics between the two infusion rates in regions near major periarterial spaces in the ventral brain surface (Figure 7). However, correlation decreased from posterior to anterior in the dorsal brain regions of inferior colliculi, superior colliculi, and neocortex, with the slower rate showed faster transport kinetics. The proximity of superior and inferior colliculi to the superior sagittal sinus suggests several potential CSF flow routes that may explain these observations, including reabsorption through arachnoid villi and meningeal lymphatic vessels, or the influence of complex dural and meningeal structures. However, the spatial resolution of MRI was insufficient to identify the structures involved in solute transport in these regions. Future research could employ advanced imaging techniques, such as simultaneous MRI and two-photon fluorescence microscopy, to better delineate the CSF flow pathways ^44^.

**Figure 7.**
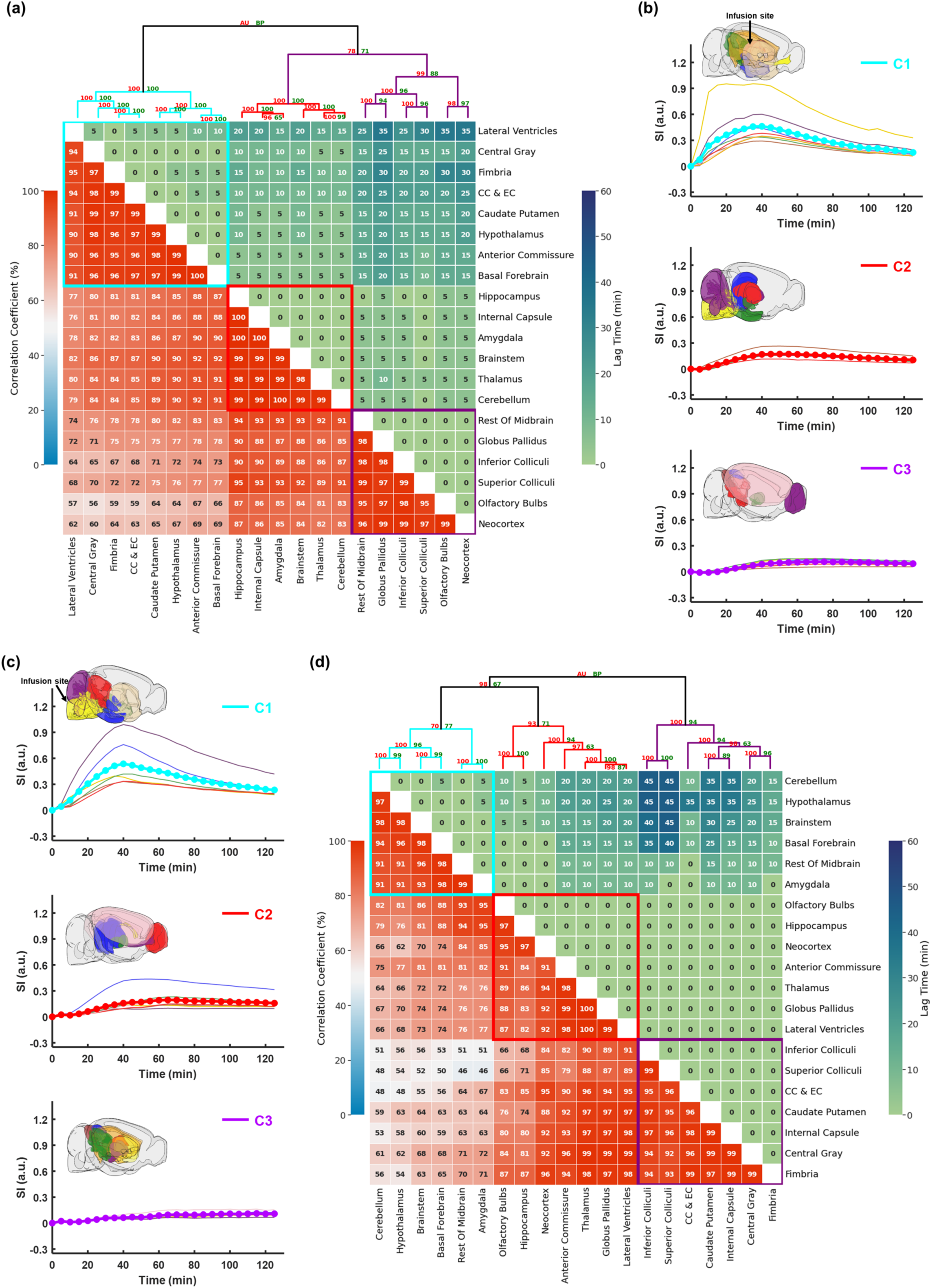
Correlation-matrix-based clustering analysis of Gd-DTPA transport in ICV-K/X (ICV, a-b) and ICM-K/X (ICM, c-d) mice. (a&d) Matrix of maximal cross-correlation (lower left) and lag time (upper right), and cluster dendrogram with bootstrap analysis (top). Values at nodes are the approximately unbiased (AU) p-values (red, left) and the bootstrap probability (BP) values (green, right), respectively. Clusters with a dissimilarity value of < 0.4 are indicated by the rectangles. (b-c) Time courses of signal changes in each cluster (thick lines). Thin lines represent signal changes in individual brain regions within the same cluster. Outlines of brain regions in each cluster are illustrated.

## Conclusion

This study used DCE-MRI to investigate the impact of infusion and anesthesia conditions on Gd-DTPA transport in mouse brain. Detailed analyses included region- specific time-course analysis of Gd-induced signal changes, and cross-correlation-based hierarchical clustering. Our findings demonstrated that K/X anesthesia promoted periarterial transport compared to isoflurane. Furthermore, the observed differences in Gd-DTPA transport patterns and kinetics regarding different infusion sites indicate the role of the ventricular-parenchyma interface in mediating CSF transport.

## Data availability

Experimental data, images, and code from this study are available upon request to the corresponding author.

## Author contributions

**Yuran Zhu:** conceptualization, methodology, investigation, software, formal analysis, visualization, writing – original draft, and funding acquisition. **Junqing Zhu:** methodology, investigation; **Chenxin Ni:** formal analysis; **Anbang Chen:** formal analysis; **Longshun Li:** methodology; **Yue Gao:** methodology; **Andrew J. Shoffstall:** methodology. **Xin Yu:** conceptualization, methodology, investigation, formal analysis, writing - original draft, funding acquisition, and project administration. **All authors:** writing - review & editing.

## Acknowledgments

This work was supported by grants from the National Institute of Health awards R01 NS124206 to X. Y., and predoctoral fellowship award from American Heart Association 23PRE1017924 to Y. Z.. The authors would like to thank Drs. Xiaoqing Alice Zhou and Xin Yu for their assistance in establishing the intracerebroventricular infusion protocol.

## Supplementary material

Supplemental material for this article is available online.

## Supplementary figures

**Fig. S1.**
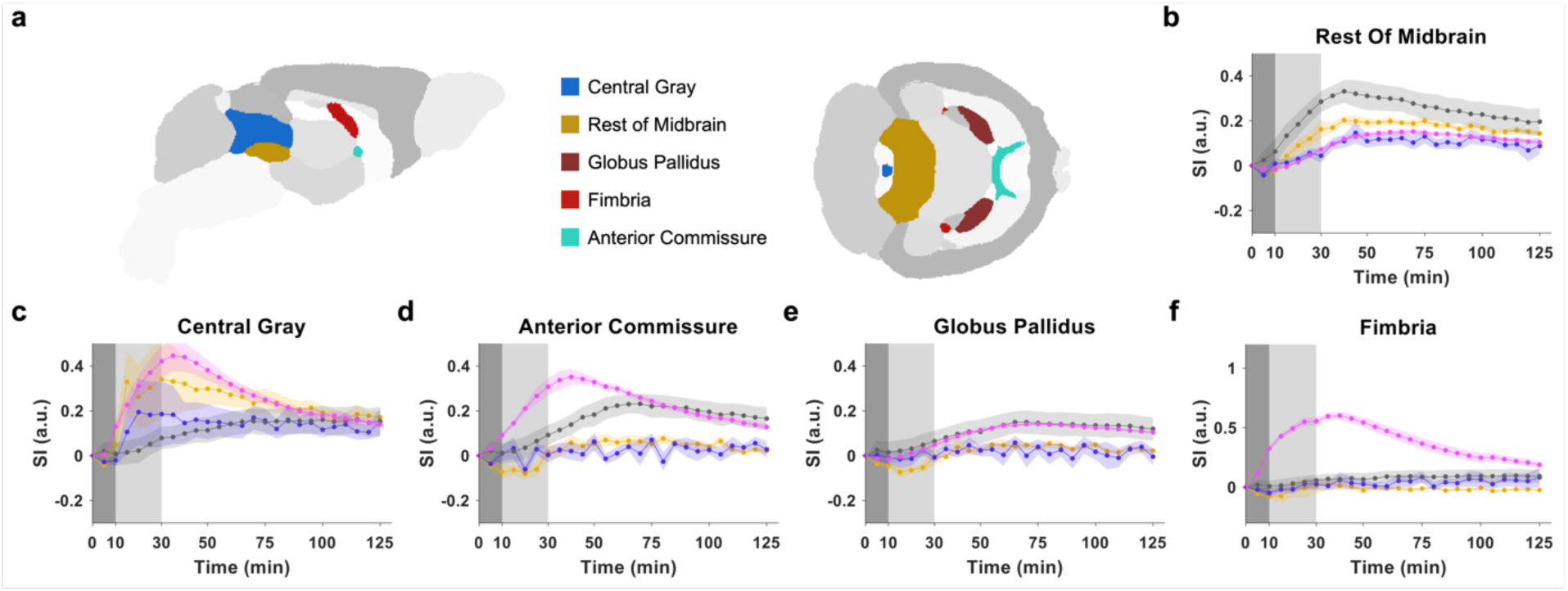
Dynamics of contrast agent transport in additional brain regions. (a) Segmentation of selected ROIs. (b)-(f) Time courses of signal changes in the selected ROIs. Gray bands indicate the time period of contrast agent infusion. Gray, magenta, yellow, and blue lines represent the mean time courses of signal changes. Shaded areas represent standard errors. ICM, intra-cisterna-magna infusion; ICV, intracerebroventricular infusion; ISO, isoflurane anesthesia; K/X, ketamine/xylazine anesthesia.

